# Hypomyelination reduces parvalbumin interneuron density and auditory cortex inhibitory function

**DOI:** 10.1101/2020.06.23.167833

**Authors:** Beatriz de Carvalho Borges, Xiangying Meng, Patrick Long, Patrick Oliver Kanold, Gabriel Corfas

## Abstract

For a long time, myelin was thought to be restricted to excitatory neurons, and studies on dysmyelination focused primarily on excitatory cells. Recent evidence showed that axons of inhibitory neurons in the neocortex are also myelinated, but the role of myelin on inhibitory circuits remains unknown. Here we studied the impact of mild hypomyelination on both excitatory and inhibitory connectivity in the primary auditory cortex (A1) with well-characterized mouse models of hypomyelination due to loss of oligodendrocyte ErbB receptor signaling. Using laser-scanning photostimulation, we found that mice with mild hypomyelination have reduced functional inhibitory connections to A1 L2/3 neurons without changes in excitatory connections, resulting in altered excitatory/inhibitory balance. These effects are not associated with altered expression of GABAergic and glutamatergic synaptic components, but with reduced density of parvalbumin-positive (PV^+^) neurons, which reflects reduced PV expression by interneurons rather than PV^+^ neuronal loss. While immunostaining shows that hypomyelination occurs in both PV^+^ and PV^-^ axons, there is a strong correlation between MBP and PV expression suggesting that myelination influences PV expression. Together, the results demonstrate that mild hypomyelination impacts A1 neuronal networks, reducing inhibitory activity, and shifting networks towards excitation.

## INTRODUCTION

It has long been known that myelination increases conduction velocity of action potentials and provides metabolic support to axons (Foster, Bujalka, & Emery, 2019; Monje, 2018). Therefore, myelination is critical for the normal patterns of neural circuits’ activation and synchronization, as well as for normal physiological, cognitive, and behavioral performance. Given the abundant presence of myelin in the mammalian central nervous system, it’s disruption can cause severe information processing defects, and has been associated with neurodegenerative and psychiatric disorders (Du & Ongur, 2013; Gibson, Geraghty, & Monje, 2018; Meuth et al., 2010).

Whereas myelin was thought to be restricted primarily to excitatory neurons, it has recently been appreciated that in the neocortex, myelin is also present on inhibitory neurons (Micheva et al., 2016). EM-based studies indicate that axon collaterals of inhibitory neurons, especially those of a large proportion of parvalbumin-positive (PV^+^) cells, are myelinated (Micheva et al., 2016; Peters & Proskauer, 1980; Tamas, Buhl, & Somogyi, 1997). PV^+^ cell axons constitute about half of all myelinated axons in somatosensory cortical layer 2/3 (L2/3), and a quarter of myelinated axons in L4, and myelination of GABAergic axons is enriched in MBP expression (Micheva et al., 2016). However, the impact of myelin defects on inhibitory circuits and neurons remains unknown.

Most insights into the impact of myelin on the function of neural circuits have been obtained by analyzing the consequences of severe demyelination, which can affect regions throughout the brain, including the cerebral cortex. For example, cuprizone-induced demyelination promotes hyper- and depolarizing shifts of the resting membrane potential of auditory thalamocortical pathway neurons and reduction in action potential firing frequency of primary auditory cortex (A1) neurons (Ghaffarian et al., 2016). Furthermore, focal demyelination in A1 permanently disrupts its tonotopic organization (Cerina et al., 2017) and auditory frequency-specific responses in the medial geniculate body (Narayanan et al., 2018). Whether these effects were mediated by myelin in excitatory or inhibitory neurons is unknown.

Moreover, in recent years there has been an increased appreciation that myelination is not an all-or-none process, but rather a continuum based on subtle differences in myelin thickness of the density of myelin segments (Monje, 2018). Furthermore, we now know that central nervous system (CNS) myelin thickness and density is influenced by experience, i.e. myelin is negatively influenced by deprivation (Liu et al., 2012; Makinodan, Rosen, Ito, & Corfas, 2012) and increased by neuronal activity and novelty (Gibson et al., 2014; McKenzie et al., 2014). Additionally, a recent study showed that while both excitatory and PV^+^ inhibitory neurons undergo homeostatic myelin remodeling under normal vision, monocular deprivation only induces adaptive myelin remodeling in PV^+^ interneurons (Yang, Michel, Jokhi, Nedivi, & Arlotta, 2020). However, much less is known about how subtle changes in myelin alters excitatory/inhibitory balance.

To gain insights into the impact of hypomyelination on inhibitory and excitatory neuronal network function, we studied A1 in mice with mild hypomyelination caused by loss of oligodendrocyte ErbB receptor signaling, either by oligodendrocyte specific expression of dominant-negative ErbB4 (Roy et al., 2007), or oligodendrocyte-specific inducible ErbB3 knock-out (Makinodan et al., 2012). To evaluate whether hypomyelination alters intracortical neural circuits in A1, we used laser-scanning photostimulation (LSPS) to optically probe functional intracortical circuits to L2/3 neurons. We found that hypomyelination leads to a reduction in inhibitory but not excitatory circuits and a consequent alteration in excitatory/inhibitory balance in L2/3 neurons. The functional alterations are not associated with changes in gene expression of molecules involved in GABAergic and glutamatergic synaptic function, but rather to a reduced density of PV^+^ neurons and lower number of myelinated PV^+^ axons in A1. Remarkably, the reduced number of PV^+^ neurons is not due to neuronal loss, but rather to a reduction in PV expression by interneurons. These results show that subtle defects in myelination can lead to large changes in gene expression and function of PV interneurons which result in large-scale changes in network function in the neocortex.

## MATERIALS AND METHODS

### Animals

All animal procedures were carried out with prior approval from the University of Michigan and the University of Maryland Committees on Use and Care of Animals in accordance with the National Research Council Guide for the Care and Use of Laboratory Animals. Mice were housed in an Association for Assessment and Accreditation of Laboratory Animal Care–accredited facility in the University of Michigan. Mice were kept in a light- (12 h on/off) and temperature- (21-23°C) controlled environment and were fed with a standard chow diet (5LOD, LabDiet, USA).

Transgenic mice expressing a dominant-negative ErbB4 receptor under control of the CNPase promoter (CNP-DN-ErbB4) (Chen et al., 2006) were crossed to wild type FVB/N J mice (JAX® mice, stock # 001800). The experimental mice were those hemizygous for CNP-DN-ErbB4 or wild type littermates. To evaluate the role of NRG-1/ErbB3 signaling in the hypomyelination and expression of PV in the A1, mice expressing inducible cre recombinase under the control of the proteolipid protein promoter (PLP/creERT) were crossed to ErbB3flox/flox mice (Makinodan et al., 2012). Floxed allele recombination was induced by intraperitoneal injection of tamoxifen (Sigma, St. Louis, MO, USA) dissolved in corn oil (10 mg/ml), by the time the peripheral myelination is known to be established (from P6). For recombination between P6 and P30, mice were treated with a dose of 33 mg/kg/day. From P30 to P56, tamoxifen dose was 100 mg/kg body weight, every other day. This regimen was used because PLP expressing oligodendrocytes differentiate and mature over time until adulthood. The experimental mice were those homozygous for ErbB3 flox expressing creERT, and homozygous ErbB3 flox littermates without cre recombinase expression. All mice were treated with tamoxifen, regardless of genotype. Experiments were performed in 2-month age mice.

The genotypes of mice were confirmed by PCR detection of the transgenes in tail-derived DNA from the CNP-DN-ErbB4, wild type, PLP/creERT and ErbB3fl/fl mice at weaning and at the end of experiments. The following pairs of primers were used: CNP-DN-ErbB4: F: 5’ TGCTGAAGGAATGGTGTGC 3’; R: 5’ CTTGTCGTCATCGTCTTTG 3’; PLP/CreERT: F: 5’ GATGTAGCCAGCAGCATGTC 3; R: 5’ ACTATATCCGTAACCTGGAT 3’ and ErbB3-flox: F: 5’ CCAACCCTTCTCCTCAGATAGG 3’; R: 5’ TGTTTGTGAAATGTGGACTTTACC 3’ and R: 5’ GGCAGGCATGTTGACTTCACTTGT 3’. Expression of the DN-ErbB4 FLAG transgene was also validated by RT-qPCR in the primary auditory cortex, using the following pair of primers: F: 5’ GAGCCTTGAGAAGATTCTTG 3’; R: 5’ TGTCGTCATCGTCTTTGTAG 3’.

### Dissection of the primary auditory cortex

Male mice were anesthetized with isoflurane (Fluoriso, Vet One) and euthanized by decapitation. Brain was collected and the primary auditory cortex (A1) fragments were micro dissected (thickness: 1.0 mm) from an area located laterally to the border of the hippocampus, according to the coordinates from Paxinos and Franklin Mouse Brain Atlas (2.06 mm posterior to bregma, 3.5 mm lateral to the midline and 2.0 mm dorsoventrally). Tissues were immediately frozen in dry ice for posterior processing for RT-qPCR (n = 7-12/group) or Western blotting (n = 6/group) analyses. Tissue harvesting was performed between 8-10am.

### Real-time quantitative RT-PCR

Total RNA was isolated from the A1 samples using RNA extraction kit, Qiazol Reagent (RNeasy mini kit; Qiagen, Germany), and DNase treatment was performed (RNase-free; Qiagen). The complementary DNA was synthesized using iScript cDNA synthesis kit (Bio-Rad, #1708891, USA), according to the manufacturers’ protocol. Quantitative RT-PCR was performed on a CFX-96 Bio-Rad reverse transcription polymerase chain reaction detection system (Hercules, CA, USA) using iTaq Universal SYBR® Green supermix (Bio-Rad, # 172-5121, USA) and primer pairs were synthesized by IDT (Coralville, IA, USA). All samples and standard curves were run in triplicate. Water instead of complementary DNA was used as a negative control. The mRNA expression in transgenic versus control mice was determined by a comparative cycle threshold (Ct) method and relative gene copy number was calculated as normalized gene expression, defined as described previously (Stankovic & Corfas, 2003). Ribosomal protein L19 (RPL19) and GAPDH were used as housekeeping genes. The following specific oligo primers were used for the target genes: RPL19, F: 5’ACCTGGATGAGAAGGATGAG 3’; R: 5’ACCTTCAGGTACAGGCTGTG 3’; GAPDH, F: 5’ TCACTGCCACCCAGAAGA 3’; R: 5’ GACGGACACATTGGGGGTAG 3’; MBP, F: 5’ATCCAAGTACCTGGCCACAG 3’; R: 5’CCTGTCACCGCTAAAGAAGC 3’; GAD65, F: 5’ CATTGATAAGTGTTTGGAGCTAGCA 3’; R: 5’ GTGCGCAAACTAGGAGGTACAA 3’; GAD67, F: 5’ TCGATTTTTCAACCAGCTCTCTACT 3’; R: 5’ GTGCAATTTCATATGTGAACATATT 3’; VGAT, F: 5’ TCCTGGTCATCGCTTACTGTCTC 3’; R: 5’ CGTCGATGTAGAACTTCACCTTCTC 3’; GABARα1, F: 5’ CCCCGGCTTGGCAACTA 3’; R: 5’ TGGTTTTGTCTCAGGCTTGAC 3’; VGLUT1, F: 5’ TCGCTACATCATCGCCATC 3’; R: 5’ GTTGTGCTGTTGTTGACCAT 3’; VGLUT2, F: 5’ CTGCGATACTGCTCACCTCTA 3’; R: 5’ GCCAACCTACTCCTCTCCAA 3’; GRIA1, F: 5’ GCTATTCCTACCGACTTGA 3’; R: 5’ CCACATCTGCTCTTCCATA 3’; GRIA2, F: 5’ CCTCATCATCATCTCCTCCTAC 3’; R: 5’ GAGCCAGAGTCTAATGTTCCA 3’; GRIA3, F: 5’ TCTAAGCCTGAGCAATGTG 3’; R: 5’ CCTTCTCTGTATGTAGCGTAAT 3’ and GRIA4, F: 5’ GCATACCTTGACCTCCTTCTG 3’; R: 5’ GCACGAACTGGCTCTCTC 3’, PARVALBUMIN: F: 5’ GCAAGATTGGGGTTGAAGAA 3’; R: 5’ GTGTCCGATTGGTACAGCCT 3’, VIP: F: 5’ CTGGCCTCTCTTTGGACCAC 3’; R: 5’ ACGGCATCAGAGTGTCGTTT 3’, SOMATOSTATIN: F: 5’ CCAACTCGAACCCAGCAATG 3’; R: 5’ TCAGAGGTCTGGCTAGGACA 3’ and ErbB3: F: 5’ TTGCCTACAGGAACGCTTACCCG 3’; R: 5’ ACCCCCCAAAACCGCAGAATC 3”. Changes in mRNA expression were calculated as relative expression (arbitrary units) respective to the wild-type group.

### Western blot analysis

Total protein from A1 was extracted using RIPA buffer (#R0278, Sigma Aldrich, USA) and protease inhibitor cocktail kit (#78410, ThermoFisher Scientific, USA). Homogenates were centrifuged at 4°C and 14,000 g for 15 minutes. Aliquots of the lysates containing 10 mg of protein were denatured in Laemmli buffer and β-mercaptoethanol (Bio-Rad, USA) at 95°C for 5 min. After electrophoresis (Mini-protean TGX gel, #456-1086, Bio-Rad, USA), samples were blotted onto nitrocellulose membranes (Immobilon-PSQ, #ISEQ00010, Merk Millipore, USA). Nonspecific binding was prevented by immersing the membranes in blocking buffer (5% BSA in Tris-buffered saline -Tween 20, TBS-T) for 60 minutes at room temperature. The membranes were then exposed overnight to the primary antibodies: mouse anti-GAPDH (1:3000, MA5-15738, RRID: AB_10977387, ThermoFisher Scientific, USA) and rat anti-MBP (1: 1,000, MAB386, RRID: AB_94975, Millipore, Germany). The blots were rinsed in TBS-T and then incubated with horseradish peroxidase-conjugated anti-mouse antibody (1:4,000, SC-516102, RRID: AB_2687626, Santa Cruz, USA) or anti-rat antibody (1: 4,000, #7077, RRID: AB_10694715, Cell Signaling, USA) for one hour at room temperature. Antibody-antigen complexes were visualized by detecting enhanced chemiluminescence using a Pierce ECL detection system (#32209, ThermoFisher Scientific, USA) and digital images with Chemi Doc Touch Image System (Bio-Rad, USA). Expression of MBP was normalized to the expression of GAPDH. Data were analyzed as relative expressions (arbitrary units) respective to wild type group.

### Immunostaining and image processing

To assess the pattern of expression of the MBP (4 – 8/group) or PV (8 – 10/group) in A1, PFC and M1 CNP-DN-ErbB4, PLP/creERT:ErbB3fl/fl or their respective littermate controls (wild types or ErbB3fl/fl mice), were anesthetized using isoflurane (Fluriso, Vet One) and transcardially perfused with saline, followed by 4% formaldehyde in 0.1 M phosphate buffer (PBS). Brains were dissected, post-fixed in the same fixative for 1 hour, placed in PBS containing 30% sucrose and sectioned on a cryostat (30-mm sections, 4 series) in the frontal plane. For MBP staining, brain coronal sections were rinsed with PBS and nonspecific binding was prevented by immersing the sections in blocking buffer (PBS, 5% normal horse serum and 0.3% Triton X-100) for one hour at room temperature. The sections were incubated overnight at 37°C with primary antibody rat anti-MBP (1:1,000, MAB386, RRID: AB_94975, Millipore, Germany) in PBS, 1% normal horse and 0.3% Triton X-100 solution. After rinses, sections were incubated for one hour with the chicken anti-rat AlexaFluor 647 secondary antibody (1:400, #A-21472, RRID: AB_2535875, Thermo Fisher Scientific, USA). For PV staining, brain coronal sections were rinsed with TBS and nonspecific binding was prevented by immersing the sections in blocking buffer (TBS, 5% normal goat serum and 0.2% Triton X-100) for one hour at room temperature. The sections were incubated overnight at 4°C with primary antibody rabbit anti-PV (1:1,000, PV25, RRID: AB_10000344, Swant). After rinses, sections were incubated for one hour with the goat anti-rabbit AlexaFluor 488 secondary antibody (1:400, #A-11094, RRID: AB_221544, Thermo Fisher Scientific, USA). Finally, the sections were mounted on superfrost microscope slides (Fisher Scientific, USA) and coversliped with Fluoro-Gel II with DAPI mounting medium (#17985-50, Electron Microscopy Sciences, USA). For double-labeling, tissue has been stained for PV, followed by MBP, as described above. For PV and Vicia Villosa Agglutinin (VVA) co-staining, mice (n = 6 – 7) were perfused as described above. PV staining was performed as above and then sections were incubated at 4°C overnight with byotinilated VVA (1:5000, #B-1235, RRID: AB_2336855, Vector Laboratories, USA). After rinsing, sections were incubated for an additional two hours with Alexa 594 streptavidin-conjugated antibody (1:500, #S32356, RRID: N/A, Thermo fisher Scientific, USA). Sections were imaged after rinsing. Quantification of PV^-^VVA cells has been performed in 3 – 4 A1 sections/mouse.

Series of systematically selected brain sections (30-mm thick, every 120-mm) representing the A1 starting on 2.06 mm posterior to bregma and ending on bregma – 3.5), the PFC (starting on 2.22 mm anterior to bregma and ending on 1.54 mm anterior to bregma) and the M1 (starting on 2.15 mm anterior to bregma and ending on 1.98 mm anterior to bregma) were acquired using a Leica SP8 confocal laser scanning microscope and 10X or 63X oil-immersion lens. The immunoreactive structures were excited using lasers with the excitation and barrier filters set for the fluorochromes used (magenta for MBP, green for PV, and blue for DAPI). Quantification of PV and MBP immunofluorescence signal, area, or the number of PV positive cells were performed in 10 different fields of the A1 per mouse, and 6 different fields of the PFC and M1 per mouse, using the Image J software (FIJI version, NIH, USA). Signal intensity was calculated by converting each frame into grayscale. The area of the signal was calculated with a homogeneously adjusted threshold. Threshold value was set based on the average of autothreshold values obtained in the control group. Histograms indicating the number of pixels of PV or MBP staining within the same area of each image were recorded and expressed as Integrated Density. Averaged Integrated Density of the fields analyzed per mouse was used to compare the levels of PV and MBP expression between groups. All immunofluorescence images shown are representative of one mouse from each group.

Quantification of myelinated PV^+^ axons was performed in A1 by obtaining z-stack images (1064 × 1064 pixels) at 63X magnification (oil immersion) with 1X digital zoom at a step size of 0.5 μm. Every 5 stacks were sampled across A1, based on DAPI nuclear labeling. Immunofluorescent colocalization of MBP and PV was manually counted by a blind investigator in 5 stacks per field, 4 - 5 fields per mouse using ImageJ (FIJI). Co-localization was defined as the circumferential bordering of a PV-labeled axon by an MBP^+^ myelin signal (Stedehouder et al., 2019; Stedehouder et al., 2017).

To estimate the area of total MBP, non-PV associated MBP, and PV associated MBP signal, we performed immunostaining for PV and MBP and image acquisition as described above. Using the FIJI software, we generated a mask for the PV signal and subtracted it from the double labeled image, thus removing all the MBP signaling that would be associated with the PV. Quantification of the area covered by total MBP, by non-PV associated MBP and by PV associated MBP in A1 was performed in 5 images/mouse, in a 63X magnification.

### In vitro Laser-Scanning Photostimulation (LSPS)

LSPS experiments were performed as previously described (Meng, Kao, Lee, & Kanold, 2015; Meng et al., 2020; Meng, Winkowski, Kao, & Kanold, 2017).

### Slice preparation

Mice were deeply anesthetized with isoflurane (Halocarbon). A block of brain containing the A1 and the medial geniculate nucleus (MGN) was removed and thalamocortical slices (500 μm thick) were cut on a vibrating microtome (Leica) in ice-cold ACSF containing (in mM): 130 NaCl, 3 KCl, 1.25 KH_2_PO_4_, 20 NaHCO_3_, 10 glucose, 1.3 MgSO_4_, 2.5 CaCl_2_ (pH 7.35 – 7.4, in 95%O_2_-5%CO_2_). The cutting angle was ~15 degrees from the horizontal plane (lateral raises) and A1 was identified as described previously (Meng et al., 2015; Meng et al., 2020; Meng et al., 2017). Slices were incubated for 1 hr in ACSF at 30°C and then kept at room temperature. Slices were held in a chamber on a fixed-stage microscope (Olympus BX51) for recording and superfused (2-4 ml/min) with high-Mg2+ ACSF recording solution at room temperature to reduce spontaneous activity in the slice. The recording solution contained (in mM): 124 NaCl, 5 KCl, 1.23 NaH_2_PO_4_, 26 NaHCO_3_, 10 glucose, 4 MgCl_2_, 4 CaCl_2_. The location of the recording site in A1 was identified by landmarks (Meng et al., 2015; Meng et al., 2020; Meng et al., 2017).

### Electrophysiology

Whole-cell recordings from L2/3 cells were performed with a patch clamp amplifier (Multiclamp 700B, Axon Instruments) using pipettes with input resistance of 4 – 9 MΩ. Data acquisition was performed with National Instruments AD boards and custom software (Ephus) (Suter et al., 2010), which was written in MATLAB (Mathworks) and adapted to our setup. Voltages were corrected for an estimated junction potential of 10 mV. Electrodes were filled with (in mM): 115 cesium methanesulfonate (CsCH_3_SO_3_), 5 NaF, 10 EGTA, 10 HEPES, 15 CsCl, 3.5 MgATP, 3 QX-314 (pH 7.25, 300 mOsm). Cesium and QX314 block most intrinsic active conductances and thus make the cells electrotonically compact. Series resistances were typically 20-25 MΩ. Photostimulation: 0.5 – 1 mM caged glutamate (*N*-(6-nitro-7-coumarinylmethyl)-L-glutamate; Ncm-Glu) (Muralidharan et al., 2016) is added to the ACSF. This compound has no effect on neuronal activity without UV light (Muralidharan et al., 2016). UV laser light (500 mW, 355 nm, 1 ms pulses, 100 kHz repetition rate, DPSS) was split by a 33% beam splitter (CVI Melles Griot), attenuated by a Pockels cell (Conoptics), gated with a laser shutter (NM Laser), and coupled into a microscope via scan mirrors (Cambridge Technology) and a dichroic mirror. The laser beam in LSPS enters the slice axially through the objective (Olympus 10·, 0.3NA/water) and has a diameter of < 20 μm. Laser power at the sample is < 25 mW. Laser power was constant between slices and recording days. We typically stimulated up to 30 · 25 sites spaced 40 μm apart, enabling us to probe areas of 1 mm2; such dense sampling reduces the influence of potential spontaneous events. Repeated stimulation yielded essentially identical maps. Stimuli were applied at 1 Hz. Analysis was performed essentially as described previously with custom software written in MATLAB (Meng et al., 2015; Meng et al., 2020; Meng et al., 2017). To detect monosynaptically evoked postsynaptic currents (PSCs), we detected PSCs with onsets in an approximately 50-ms window after the stimulation. This window was chosen based on the observed spiking latency under our recording conditions (Meng et al., 2015; Meng et al., 2020; Meng et al., 2017). Our recordings were performed at room temperature and in high-Mg2+ solution to reduce the probability of polysynaptic inputs. We measured both peak amplitude and transferred charge; transferred charge was measured by integrating the PSC. While the transferred charge might include contributions from multiple events, our prior studies showed a strong correlation between these measures (Meng et al., 2015; Meng et al., 2020; Meng et al., 2017). Traces containing a short-latency (< 8 ms) ‘direct’ response were discarded from the analysis (black patches in color-coded maps) as were traces that contained longer latency inward currents of long duration (> 50 ms) (Fig. 3b). The short-latency currents could sometimes be seen in locations surrounding (< 100 μm) areas that gave a ‘direct’ response. Occasionally some of the ‘direct’ responses contained evoked synaptic responses that we did not separate out, which leads to an underestimation of local short-range connections. Cells that did not show any large (>100 pA) direct responses were excluded from the analysis as these could be astrocytes. It is likely that the observed PSCs at each stimulus location represent the activity of multiple presynaptic cells.

Stimulus locations that showed PSC were deemed connected and we derived binary connection maps. We aligned connection maps for L2/3 cells in the population and averaged connection maps to derive a spatial connection probability map (Fig. 3c). In these maps the value at each stimulus location indicates the fraction of L2/3 cells that received input from these stimulus locations. Layer boundaries were determined from the infrared pictures. We derived laminar measures. Input area is calculated as the area within each layer that gave rise to PSCs. Mean charge is the average charge of PSCs from each stimulus location in each layer. Intralaminar integration distance is the extent in the rostro-caudal direction that encompasses connected stimulus locations in each layer. We calculated E/I balance in each layer for measures of input area and strength as (E-I)/(E+I), thus (Area_E_-Area_I_)/(Area_E_+Area_I_), resulting in a number that varied between −1 and 1 with 1 indicating dominant excitation and −1 indicating dominant inhibition.

Spatial connection probability maps show the average connection pattern in each group. To compare the large-scale connectivity between cells in each group we calculated the spatial correlation of the binary connection maps in each group by calculating the pairwise crosscorrelations (Meng et al., 2020).

### Statistical analysis

Results are expressed as means ± SEM and data were analyzed using MATLAB or GraphPad Prism 9 software. Results of qRT-PCR, Western blots and immunofluorescence experiments were analyzed by unpaired two-tailed Student’s t-test if the variables passed a normality test or by Mann-Whitney non-parametric test if not. LSPS results were analyzed by ANOVA followed by a multicomparison test. To evaluate the correlation between PV and MBP mRNA expression a Pearson’s correlation test was performed. Differences were accepted as significant at P < 0.05.

## RESULTS

### Disruption of oligodendrocyte ErbB signaling results in reduced myelin basic protein and mRNA levels in A1

We previously showed that expression of a dominant-negative ErbB4 receptor in cells of the oligodendrocyte lineage under the control of the CNPase promoter (CNP-DN-ErbB4, (Roy et al., 2007)) results in mice with reduced myelin thickness in optic nerve, corpus callosum and prefrontal cortex axons (Roy et al., 2007) as well as thinner myelin and reduced levels of MBP protein and mRNA levels in peripheral nerves (Chen et al., 2006). DN-ErbB4, which in these mice is only expressed in myelinating glia, is a truncated ErbB4 receptor, and its expression abolishes the signaling of NRG1 receptors ErbB2, 3 and 4 (Chen et al., 2006; Prevot et al., 2003; Rio, Rieff, Qi, Khurana, & Corfas, 1997) without affecting signaling by ErbB1, the receptor for EGF (Prevot et al., 2003). We now find that DN-ErbB4 expression in oligodendrocytes leads to similar alterations in A1 myelination. Quantitative immunofluorescence showed that the intensity of MBP in A1 is markedly reduced in adult mutant mice compared to their wild type (WT) littermates (Fig. 1a & b). Western blot analysis (Fig. 1c & d) showed a reduction in MBP protein in A1 of CNP-DN-ErbB4 mice, and quantitative RT-PCR showed a similar reduction in MBP mRNA levels (Fig. 1e). Furthermore, quantification of the MBP immunofluorescence density showed that the area of MBP signal is reduced in the mutant, consistent with a reduction in the number of MBP+ processes in A1 (Fig. 1f). Together, these results indicate that loss of oligodendrocyte ErbB receptor signaling leads to A1 hypomyelination.

**Figure 1:**
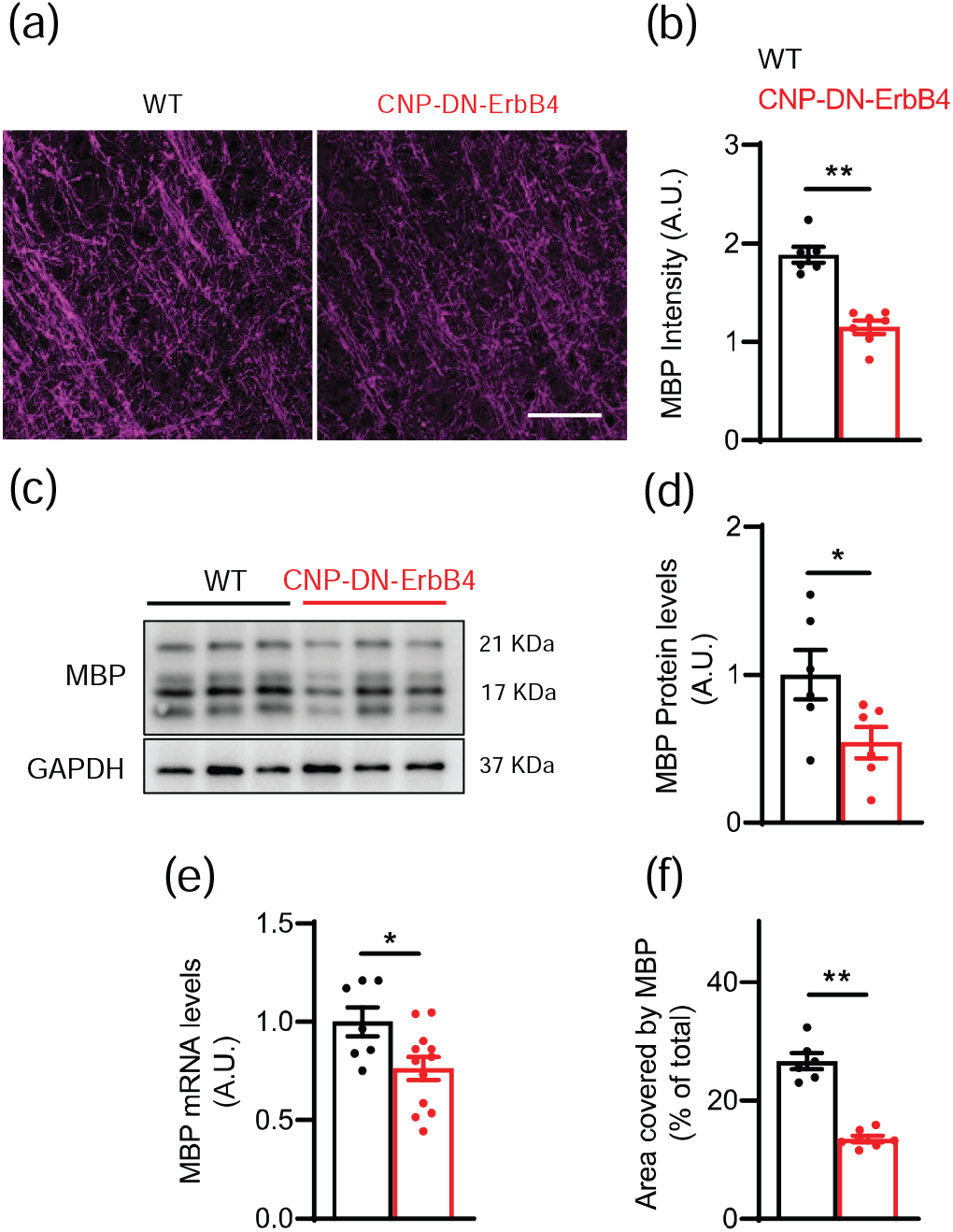
Loss of oligodendrocyte ErbB receptor signaling leads to a reduction in MBP protein and mRNA expression levels in A1. **a**, Representative photomicrographs of primary auditory cortex of WT and CNP-DN-ErbB4 mice showing the expression of myelin basic protein (MBP) (magenta). Scale bar: 20μm. **b**, Quantification of MBP staining intensity (WT: black; CNP-DN-ErbB4: red) (n = 6 – 7 mice per genotype; ** p = 0.001). A.U.: arbitrary units. **c**, Representative MBP and GAPDH Western blots of A1 samples from WT and CNP-DN-ErbB4 mice. **d**, Quantification of MBP protein expression normalized by GAPDH in A1 of WT and CNP-DN-ErbB4 mice (n = 6; p = 0.0413). **e**, MBP mRNA levels in A1 of WT and CNP-DN-ErbB4 mice (n = 7 – 12; p = 0.0226). **f**, Quantification of the percent of area covered by MBP (WT: black; CNP-DN-ErbB4: red) (n = 6 – 7 mice per genotype; ** p < 0.001). A.U.: arbitrary units. Unpaired two-tailed Student’s t test was performed. Data are expressed as mean ± SEM.

### Hypomyelination does not alter sensitivity of cortical neurons to photostimulation

We next examined the impact of hypomyelination on A1 intracortical neural circuits using laser-scanning photostimulation (LSPS) with caged glutamate (Meng et al., 2015; Meng et al., 2020; Meng et al., 2017) in brain slices of adult mice. We first tested if oligodendrocyte DN-ErbB4 expression affects the ability of A1 neurons to fire action potentials to photo released glutamate by performing cell-attached patch recordings from cells in L2/3 and L4 (N=27 L4 and N=38 L2/3 neurons in WT, N=20 L4 and N=36 L2/3 neurons in CNP-DN-ErbB4) (Fig. 2a). Short UV laser pulses (1ms) to focally release glutamate were targeted to multiple stimulus spots covering the extent of A1 (Fig. 2b). Stimulation close to the cell body led to action potentials (Fig. 2b). There were no differences between the genotypes in the area around targeted neurons where action potentials could be generated (Fig. 2b, c, L2/3: p=0.14; L4: p=0.62) or in the number of evoked action potentials (Fig. 2c, L2/3: p=0.29; L4: p=0.74). Thus, the spatial resolution of LSPS is not affected by hypomyelination.

**Figure 2:**
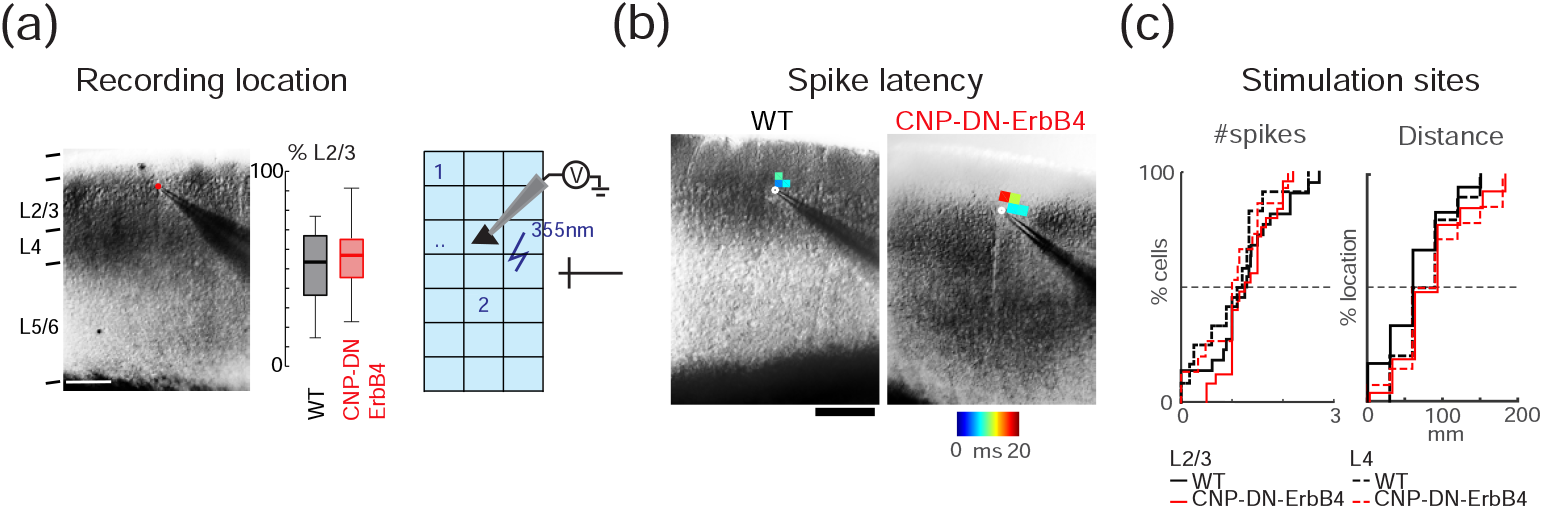
Sensitivity to photo released glutamate in Layer 2/3 neurons is normal in A1 of CNP-DN-ErbB4 mice. **a**, <u>Left</u>, Infrared image of cortical field with patch pipette on a. Scale bar is 200 μm. Cortical layers are identified based on the DIC image. <u>Right</u>, Position of recorded cells within Layer 2/3. Plotted are the relative positions within Layer 2/3 for WT (black) and CNP-DN-ErbB4 (red) mice. 0 refers to the border with Layer 4 and 100 refers to the border with Layer 1. **b**, <u>Left</u>, Graphic illustration on how cell attach LSPS experiments were performed. The cortical fields were divided into approximately 30 by 25 grids. UV laser targets stimulation sites in a grid in a pseudorandom pattern to make sure that two nearby locations won’t be stimulated sequentially to avoid adaptation. Cells under laser activation sites could be activated and generate action potentials (APs). <u>Right</u>, Cell attach recordings on Layer 2/3 and 4 cells show areas that evoke action potentials. Maps show first spike latencies encoded by color and overlaid on infrared images. **c**, Cumulative distributions of number of spikes (Left) and distances from locations that resulted in APs to the soma of L2/3 (solid) and L4 (dashed) cells for both WT (black) and CNP-DN-ErbB4 (red) mice.

### Hypomyelination does not alter excitatory connections to L2/3 neurons

To map and quantify functional excitatory inputs to L2/3 neurons we performed whole-cell recording combined with LSPS while cells were held at a holding potential of −70mV. All recorded cells were from similar laminar positions (Fig. 2a, p=0.062). We targeted the laser pulse to multiple (600-900) stimulus locations spanning across the cortical extent and recorded the resulting membrane currents (Fig. 3a). Targeting the cell body and the proximal dendrites of the cell under study caused large amplitude and short-latency (<8ms) currents (Fig. 3b), reflecting direct activation of glutamate receptors on the neuron. These ‘direct’ currents were excluded from the analysis. Targeting other sites could result in longer-latency (> 8ms) currents, which under our recording conditions, reflect monosynaptically evoked post-synaptic currents (PSCs) (Meng et al., 2015; Meng et al., 2020; Meng et al., 2017) (Fig. 3b).

**Figure 3:**
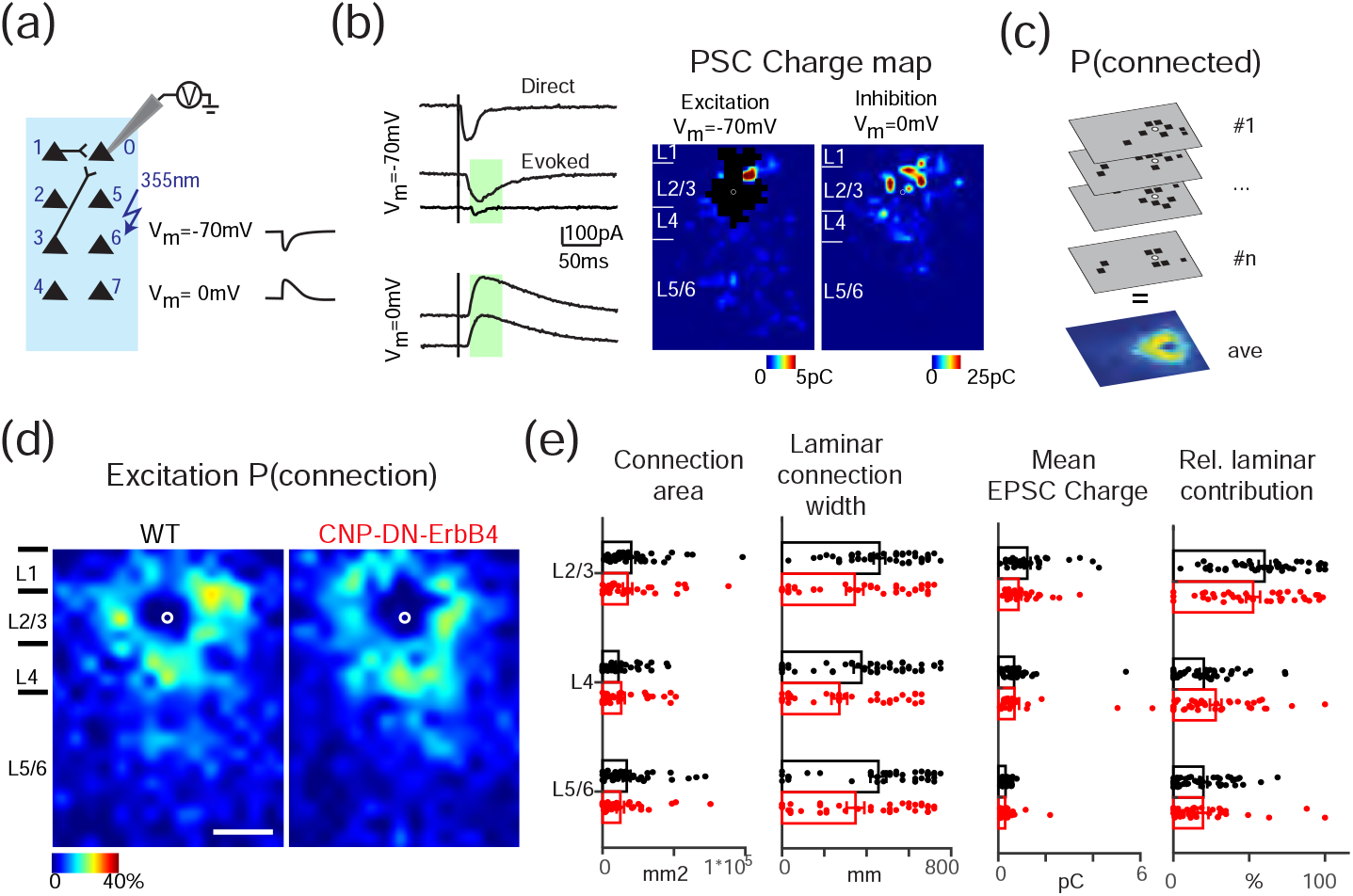
Excitatory circuits to L2/3 neurons are normal in A1 of CNP-DN-ErbB4 mice. **a**, Schematic diagram shows how whole-cell voltage-clamp combined with LSPS experiments are performed. **b**, Whole-cell voltage-clamp recordings at holding potential of −70mV (top) and 0mV (bottom). <u>Left</u>, shows the traces obtained with photostimulation at different locations. Solid lines indicate time of photostimulation. Green shaded area is the analysis window. It started at 8 ms and ended at 50ms after laser onset. <u>Right</u>, example of excitatory and inhibitory maps for a L2/3 cell. Pseudocolor indicates the PSC charge at each stimulation location. Black area indicates where direct responses are. White circle marks the soma location. Horizontal bars indicate layer borders and are 100 μm long. **c**, Cartoon illustrating how the connection probability (P(connection)) map is calculated. All the input maps contain 0 (gray, the area has no connection to the recorded cell) and 1 (black square, the area has monosynaptic connection to the recorded cell) and are aligned to soma (white circles). The P (connection) map is calculated by averaging all the input maps along the z-axis. **d**, Spatial connection probability of excitatory connections for WT (left) and CNP-DN-ErbB4 (right) mice, n=36 cells for WT and n = 32 cells for CNP-DN-ErbB4. The border bars are the averaged borders across all the cells in each group. Scale bar is 200μm long. **e**, Bar graph of total area (left) and laminar connection width (right) of excitatory inputs from L2/3, L4 and L5/6 to L2/3 neurons of WT (black) and CNP-DN-ErbB4 (red) mice. Data are expressed as mean ± SEM, p>0.05. F, Bar graph of mean charge (left) and relative laminar charge contribution (right) from L2/3, L4 and L5/6 to L2/3 neurons of WT (black) and CNP-DN-ErbB4 (red) mice. Data are expressed as mean ± SEM, p>0.05. ANOVA was performed.

We mapped 68 L2/3 cells (36 cells in 14 WT mice, 32 cells in 11 CNP-DN-ErbB4 mice) and generated spatial connection maps for each cell which indicated which stimulus location resulted in an evoked PSC. We then aligned and averaged all individual spatial connection maps to the soma position of the individual cells (Fig. 3c). By averaging the individual connection maps, we obtained the spatial connection probability map for excitatory inputs where each value (represented as color in the maps) denotes the fraction of neurons in the population that receives input from a particular location (Fig. 3d). Qualitatively, a comparison of the excitatory connection probability maps from WT and in CNP-DN-ErbB4 animals showed no differences (Fig. 3d).

Since spatial averaging can obscure differences in functional connections, we quantified the laminar changes of the connection properties for each cell as in prior studies (Meng et al., 2015; Meng et al., 2020; Meng et al., 2017). For each neuron, we quantified the total area in each layer where stimulation resulted in a response. The total area within each layer where stimulation can evoke EPSCs in L2/3 neurons was similar in all layers between genotypes (Fig. 3e, L2/3: p=0.13; L4: p=0.96; L5/6: p=0.12). The thalamocortical slices are cut such that the macroscale rostro-caudally oriented tonotopic map is preserved in the slice plane. Thus, the spatial extent of functional integration along the slice is a proxy for the integration along the tonotopic axis. To assess this laminar connection width, we calculated the distance in each layer that covers 80% of the input sites. We found that the laminar connection width is similar between genotypes indicating that the functional integration across the tonotopic axis was unchanged in CNP-DN-ErbB4 mice (Fig. 3e, L2/3: p=0.11; L4: p=0.14; L5/6: p=0.14). Circuit changes can result from both connection probability and input strength. Thus, we measured the average charge of the evoked EPSC and the relative EPSC charge contribution from each layer to L2/3 cells (Fig. 3f). Evoked EPSCs and relative contribution from all layers were not different between genotypes (Fig. 3f, evoked EPSC, L2/3: p=0.15; L4: p=0.16; L5/6: p=0.14; relative contribution, L2/3: p=0.22; L4: p=0.21; L5/6: p=0.52). Together, these results indicate that hypomyelination does not affect the spatial pattern of functional excitatory connectivity to L2/3 neurons.

### Hypomyelination leads to reduced inhibitory connectivity to L2/3 neurons

To map and quantify functional inhibitory input to L2/3 neurons we held cells at 0 mV and repeated the LSPS. Qualitative comparison of the resulting connection probability maps from WT and CNP-DN-ErbB4 animals showed an apparent reduction in inputs from L4 and from within L2/3 (Fig. 4a). We confirmed this observation by quantifying the total area within each layer where stimulation evoked IPSCs in the recorded L2/3 neurons. We found that total input from L2/3, L4 and L5/6 was reduced in CNP-DN-ErbB4 animals (Fig. 4b, L2/3: p=5.1 ×10^−3^; L4: p=4.5 ×10^−3^; L5/6: p=3.7 ×10^−3^). Moreover, the laminar connection width was smaller in all layers of CNP-DN-ErbB4 mice than in cells from WT (Fig. 4b, L2/3: p=8.5 ×10^−3^; L4: p=2.3 ×10^−3^; L5/6: p=2.9 ×10^−3^). This indicates that cells from CNP-DN-ErbB4 mice receive inputs from a more restricted area than in cells from WTs. The average charge of the evoked IPSC within L2/3 in CNP-DN-ErbB4 animals was reduced (L2/3: p=0.02; L4: p=0.23; L5/6: p=0.36) but the relative contribution from L2/3 is increased (Fig. 4c, L2/3: p=0.031; L4: p=0.11; L5/6: p=0.021). These results suggest that hypomyelination causes a spatial hypoconnectivity of inhibitory connections. As expected from the decreased inhibitory connectivity in face of unchanged excitatory connectivity, calculating the balance between excitation and inhibition showed that this balance is shifted towards excitation (Fig. 4d, EI charge: L2/3: p=0.901; L4: p=2.4 ×10^−3^; L5/6: p=0.012; EI peak: L2/3: p=0.32; L4: p=6.2 ×10^−4^; L5/6: p=6.9 ×10^−3^; EI Area: L2/3: p=0.52; L4: p=7.6 ×10^−4^; L5/6: p=0.011).

**Figure 4:**
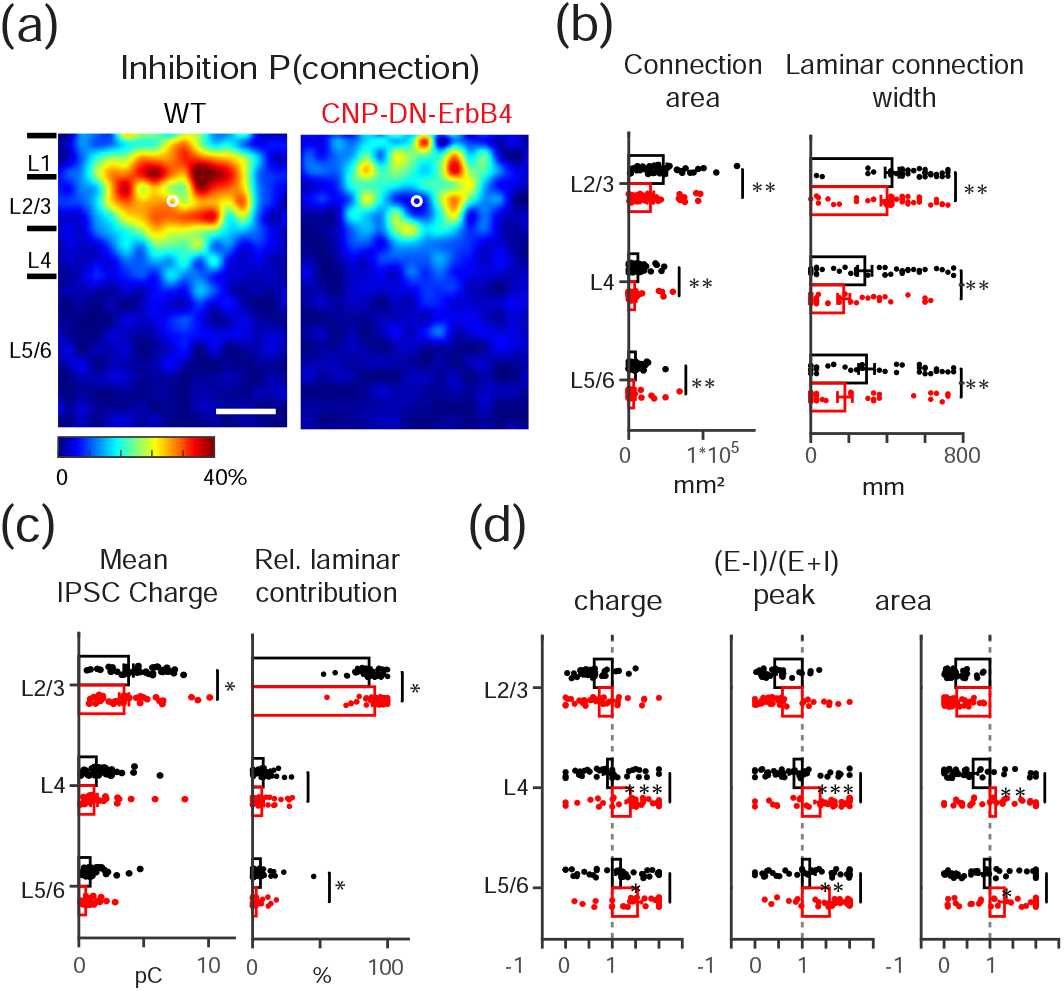
Reduced connectivity of inhibitory circuits to L2/3 neurons in A1 of CNP-DN-ErbB4 mice. **a**, Spatial connection probability of inhibitory connections for WT (left) and CNP-DN-ErbB4 (right) mice. The border bars are the averaged borders across all the cells in each group. Scale bar is 200μm long. **b**, Bar graph of total area (left) and laminar connection width (right) of inhibitory inputs from L2/3, L4 and L5/6 to L2/3 neurons of WT (black) and CNP-DN-ErbB4 (red) mice. Data are expressed as mean ± SEM. * p<0.05, **p<0.01, ***p<0.001. Loss of myelin reduces the inhibitory inputs from all layers. **c**, Bar graph of inhibitory mean charge (left) and relative laminar charge contribution (right) from L2/3, L4 and L5/6 to L2/3 neurons of WT (black) and CNP-DN-ErbB4 (red) mice. Data are expressed as mean ± SEM, *p<0.05. **d**, Excitatory/inhibitory ((E-I)/(E^+^I)) balance for inputs from L2/3, L4, and L5/6 based on charge (left), peak (middle) and area (right). L2/3 neurons in CNP-DN-ErbB4 mice receive more excitation than inhibition, * p<0.05, **p<0.01, ***p<0.001. ANOVA was performed.

Individual cells can vary in their inputs and functional circuit diversity can emerge through development (Meng et al., 2020). We thus investigated the functional spatial diversity of the circuits impinging on L2/3 cells by calculating the similarity (spatial correlation) between connection maps within each population (Meng et al., 2020). We found that the circuit similarity is increased for both excitatory and inhibitory circuits in cells from CNP-DN-ErbB4 mice (Fig. 5, Excitatory correlation: p= 3.5 ×10^−3^; Inhibitory correlation: 7.4 ×10^−24^). Thus, loss of oligodendrocyte ErbB signaling leads to a hypoconnectivity of inhibitory circuits and reorganized excitatory circuits.

**Figure 5:**
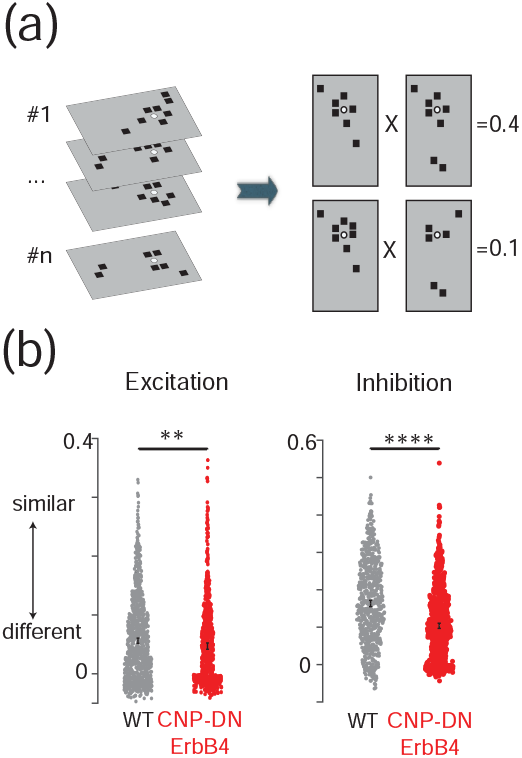
Reduction in inhibitory circuits and reorganization of excitatory circuits in A1 of CNP-DN-ErbB4 mice. **a**, Graphical representation of calculation of pairwise correlation between functional connection maps. Each black square represents the area that has monosynaptic connection to the recorded cell. Each connection map will be first vectorized and pairwise correlation between all the vectors will be calculated. **b**, Mean and SEM of pairwise correlations of both excitatory (left) and inhibitory (right) maps for WT (black) and CNP-DN-ErbB4 (red) mice. Hypomyelination reduced the pairwise correlation of both excitatory and inhibitory maps, **p<0.01, ****p<0.0001 ANOVA was performed.

### Hypomyelination does not alter expression of inhibitory or excitatory synaptic components but reduces PV mRNA levels and PV^+^ cells number in the auditory cortex

Our functional studies show changes in inhibitory circuits. Since ErbB receptor signaling has been reported to regulate neurotransmitter receptor expression in a diverse set of cells (Gerecke, Wyss, Karavanova, Buonanno, & Carroll, 2001), we tested if the levels of mRNA for several genes central to inhibitory and excitatory neurotransmission are altered in A1 in CNP-DN-ErbB4 mice. Quantitative RT-PCR analysis revealed normal mRNA levels for molecules associated with GABAergic neurotransmission (GAD65, GAD67, VGAT and GABARα1) (Fig. 6a), and glutamatergic neurotransmission (VGLUT1, VGLUT2, GRIA1, GRIA2, GRIA3 and GRIA4) (Fig. 6b). Thus, the observed circuit differences do not appear to result from alteration in expression of components of GABAergic or glutamatergic synapses.

**Figure 6:**
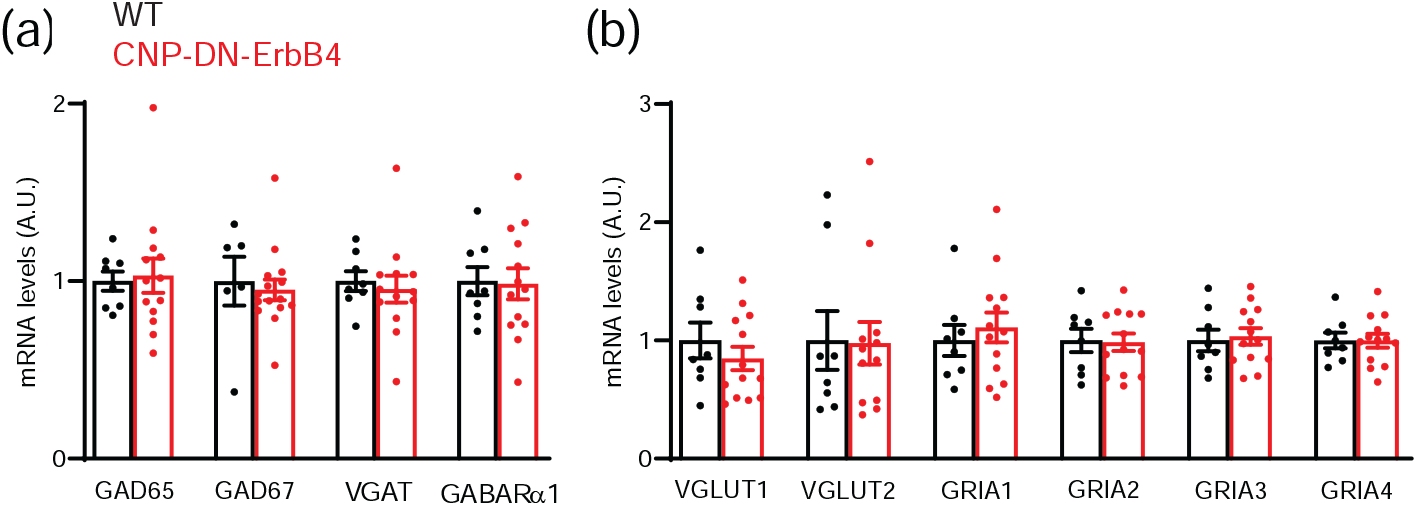
mRNA levels for excitatory and inhibitory synaptic proteins is normal in A1 of CNP-DN-ErbB4 mice. mRNA expression for synaptic proteins in A1 of WT (black) and CNP-DN-ErbB4 (red) male mice were measured by real-time RT-PCR. **a**, GABAergic markers: GAD65 (p = 0.9574), GAD67 (p = 0.8027), VGAT (p = 0.7639) and GABARα1 (p = 0.4925). **b**, Glutamatergic markers: VGLUT1 (p = 0.1843), VGLUT2 (p = 0.2377), GRIA1 (p = 0.8244), GRIA2 (p = 0.7100), GRIA3 (p = 0.5533) and GRIA4 (p = 6197). Unpaired two-tailed Student’s t test was performed. Data are expressed as mean ± SEM. No differences were found between genotypes (n = 8 – 12).

We then wondered if the observed reduction in functional connections could be due to alterations in specific types of inhibitory neurons. To test this scenario, we first measured the mRNA levels of genes that mark the three main populations of inhibitory interneurons i.e., parvalbumin (PV), vasoactive intestinal peptide (VIP) and somatostatin (SST). Only PV mRNA levels were reduced in A1 of CNP-DN-ErbB4 mice (Fig. 7a). We found a strong correlation between the levels of PV and MBP mRNA in A1 (p=0.0064) (Fig. 7b), suggesting that key aspects of PV neurons are linked to myelination. Consistent with the reduction in PV mRNA, PV immunostaining showed that the density of A1 PV^+^ neuron cell bodies is reduced in CNP-DN-ErbB4 mice (Fig. 7c,d). Moreover, intensity of PV immunostaining in the neuropil is also lower, suggesting that the number of PV^+^ axons is reduced as well (Fig. 7e-h).

**Figure 7:**
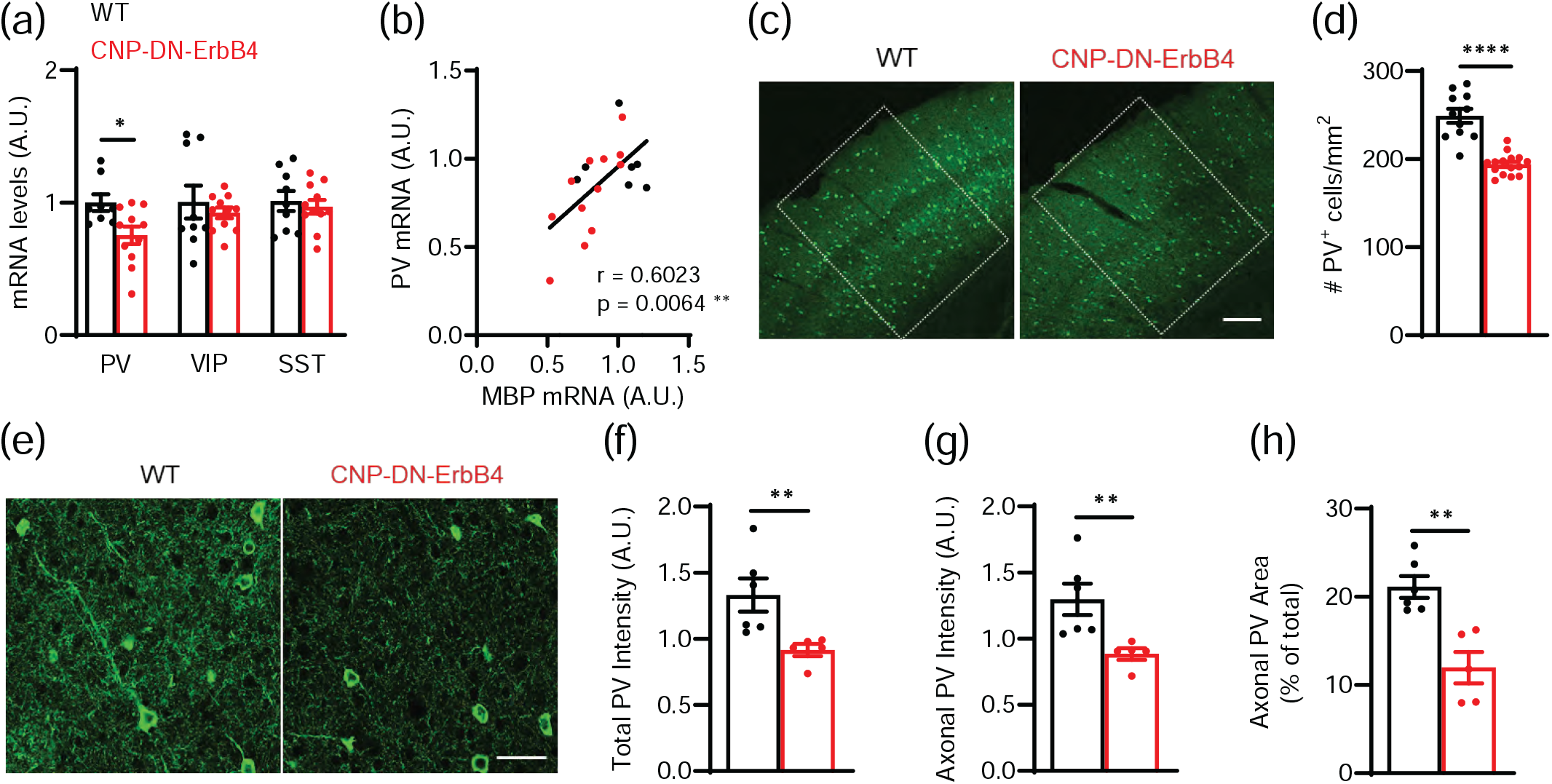
Loss of oligodendrocyte ErbB signaling leads to reduced A1 PV^+^ cell density and PV mRNA levels. **a**, mRNA levels for parvalbumin (PV), vasoactive intestinal peptide (VIP) and somatostatin (SST) in the A1 of WT (black) and CNP-DN-ErbB4 (red) mice. PV mRNA levels are reduced in A1 of CNP-DN-ErbB4 mice (n = 8 – 11; p = 0.0290) whereas no changes are observed in VIP (p = 0.51) and STT (p = 0.6301) mRNA. **b**, PV mRNA levels correlate with MBP mRNA levels in A1 (WT: black dots; CNP-DN-ErbB4: red dots; Pearson’s correlation r = 0.6023; p = 0.0064) **c**, Representative photomicrographs of A1 sections from WT and CNP-DN-ErbB4 mice showing PV^+^ cells (green). Scale bar: 100μm. **d**, A1 PV^+^ cell density is reduced in CNP-DN-ErbB4 mice (n = 10 – 14; p <0.0001). **e**, Representative photomicrographs of A1 sections from WT and CNP-DN-ErbB4 mice showing PV^+^ neurons and axons (green). Scale bar: 20μm. **f**, Quantification of total PV staining intensity (cell bodies + axons) in WT (black) and CNP-DN-ErbB4 (red) mice (n = 5 – 6, p = 0.0043). A.U.: arbitrary units. **g**, Quantification of axonal PV staining intensity (total PV intensity – cell body PV intensity) in WT (black) and CNP-DN-ErbB4 (red) mice (n = 5 – 6, p = 0.0043). A.U.: arbitrary units. **h**, Quantification of the area covered by PV^+^ axons in WT (black) and CNP-DN-ErbB4 (red) mice (n = 5 – 6; p = 0.0087). A.U.: arbitrary units. Unpaired two-tailed Student’s t test or Mann-Whitney test were performed. Data are expressed as mean ± SEM.

To determine if the reduced A1 PV^+^ neuron density in CNP-DN-ERrB4 mice is due to neuronal loss or altered PV expression, we co-labeled A1 sections with PV antibodies and Vicia Villosa Agglutinin (VVA), a lectin that preferentially binds to the extracellular matrix enwrapping PV neurons (Drake, Mulligan, Wimpey, Hendrickson, & Chavkin, 1991; Luth, Fischer, & Celio, 1992) (Fig 8a). PV^+^ and VVA^+^ cells were identified by immunofluorescence and confocal microscopy. Quantitative analysis showed that, whereas PV^+^ neuron density is significantly reduced in CNP-DN-ErbB4 mice, VVA^+^ neuron density remains unchanged (Fig 8b, c). Importantly, there was a clear impact on the pattern of PV and VVA double labeled neurons. While 84% of VVA^+^ cells also expressed PV in controls, only 67% did in CNP-DN-ErbB4 mice (Fig 8d). In contrast, the proportion of PV^+^ cells that were also VVA^+^ was not altered by the hypomyelination (87.8% in controls and 86% in CNP-DN-ErbB4 mice) (Fig 8e). These results show that the reduction in A1 PV^+^ neurons is due to reduced PV expression by some interneurons rather than their loss. Together, these results suggest that in A1 hypomyelination affects PV neurons, and that myelination, PV expression and PV^+^ neuron abundance are linked.

**Figure 8:**
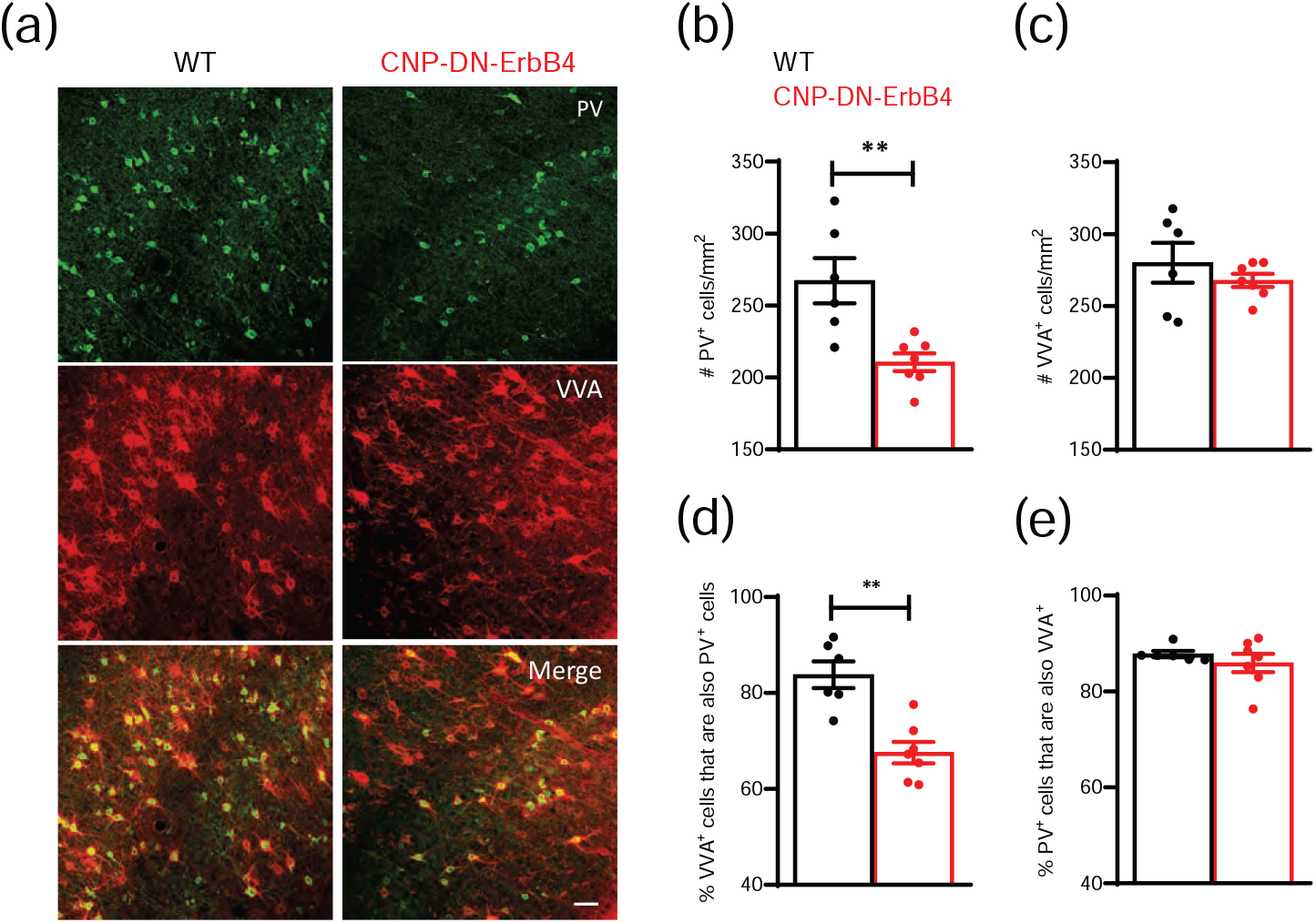
Hypomyelination leads to reduced density of PV^+^ neurons but normal density of VVA^+^ cells in A1. **a**, Representative photomicrographs of A1 sections from WT and CNP-DN-ErbB4 mice showing PV^+^ (green) and VVA^+^ (red) immunostaining. Scale bar: 50μm. **b**, A1 PV^+^ cell density is reduced in CNP-DN-ErbB4 mice (n = 6 – 7; p = 0.0058). **c**, A1 VVA^+^ cell density is not altered in CNP-DN-ErbB4 mice (n = 6 – 7; p = 0.60). **d**, The fraction of VVA^+^ neurons also expressing PV is reduced in CNP-DN-erbB4 mice (67.5%) in comparison to WT (84%) (n = 6 – 7; p = 0.0023). **e**, The fraction of PV^+^ neurons also expressing VVA is similar between groups (WT: 87.8%; CNP-DN-ErbB4: 86% (n = 6 – 7; p = 0.73). Mann-Whitney test was performed. Data are expressed as mean ± SEM.

### Loss of oligodendrocyte ErbB signaling due to expression of DN-ErbB4 reduces the number of myelinated PV^+^ axons in the A1

To determine if A1 PV^+^ interneurons are hypomyelinated in CNP-DN-ErbB4, we quantified the density of PV^+^ axonal segments that are ensheated by MBP using immunofluorescence and confocal imaging as done previously by Stedehouder et al. (Stedehouder et al., 2019; Stedehouder et al., 2017) (Fig. 9a,b). We found a 57% reduction in the number of PV^+^/MBP^+^ axonal segments in A1 of CNP-DN-ErbB4 mice (Fig. 9a,c). Furthermore, we observed a strong correlation between the intensity of PV and MBP signal in A1 axons (r = 0.6021, p,0.0001) (Fig. 9d), and between the area covered by PV^+^ and MBP^+^ in the neuropil (r = 0.6373, p<0.0001) (Fig. 9e), supporting the notion that myelination of axons of PV^+^ neurons are negatively impacted by disruption of ErbB receptor signaling and might explain the changes in E/I balance in the A1 of CNP-DN-ErbB4 mice.

**Figure 9:**
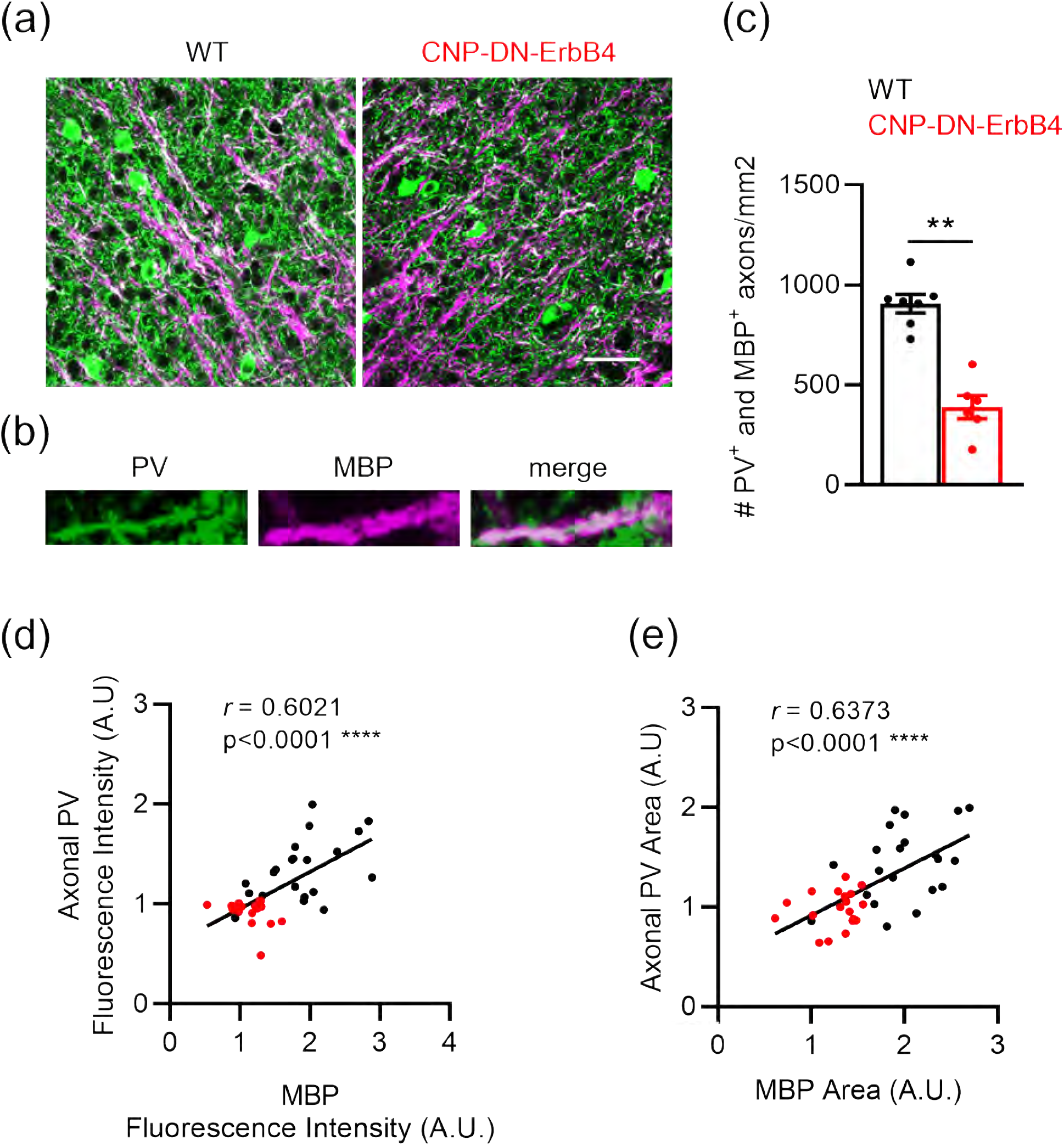
Loss of oligodendrocyte ErbB signaling leads to reduced number of A1 PV^+^ axons associated with MBP staining, reduced intensity of PV axonal immunostaining, and the area of PV^+^ axons. The intensity and area of PV^+^ axons correlate with the intensity and area of their associated MBP immunostaining. **a**, Representative photomicrographs of A1 sections from WT and CNP-DN-ErbB4 mice showing PV^+^ (green) and MBP^+^ (magenta) immunostaining. Scale bar: 20μm. **b**, Representative photomicrographs of 2X zoom images from the 63X images shown in panel **c**, Density of PV-MBP double-labelled axons is reduced in A1 of CNP-DN-ErbB4 mice (n = 6 – 7; p = 0.0012). **d**, A1 axonal PV fluorescence intensity correlates with MBP fluorescence intensity (WT: black dots, CNP-DN-erbB4: red dots; Pearson’s correlation r = 0.6021; p <0.0001).. **e**, A1 axonal PV area correlates with MBP area (WT: black dots, CNP-DN-erbB4: red dots; Pearson’s correlation r = 0.6373; p <0.0001). Mann-Whitney test was performed. Data are expressed as mean ± SEM.

### Loss of oligodendrocyte ErbB signaling leads to reduction in MBP associated with both PV^+^ and PV^-^ axons

Since cortical myelin is present on axons of both excitatory neurons and PV^+^ interneurons (Micheva et al., 2016; Peters & Proskauer, 1980; Tamas et al., 1997), it was important to determine if loss of oligodendrocyte ErbB signaling affects these axons differentially. To this end, we used quantitative immunofluorescence on a new set of samples (Fig. 10). As we found earlier (Fig. 1), the area covered by MBP immunoreactivity was reduced in A1 of CNP-DN-ErbB4 mice (Fig. 10a & b). To identify the portion of the image reflecting MBP associated with PV^-^ axons, we created a mask for all the PV^+^ pixels and subtracted it from the MBP image (Fig. 10a). The area of MBP immunofluorescence associated with PV^+^ axons was calculated as the difference between total MBP area and the MBP not associated with PV signal. Both portions of the MBP signal were reduced in CNP-DN-ErbB4 (Figs. 10c and d), suggesting that ErbB receptor signaling regulates myelination of both PV^+^ and PV^-^ axons, the latter most probably representing axons from excitatory neurons. The area of MBP associated with PV^+^ pixels was reduced to a larger extent, most probably reflecting the reduced abundance of PV^+^ axons in CNP-DN-ErbB4. However, we cannot rule out the possibility that myelination of PV^+^ axons in A1 has a stronger dependence on ErbB signaling than excitatory axons.

**Figure 10:**
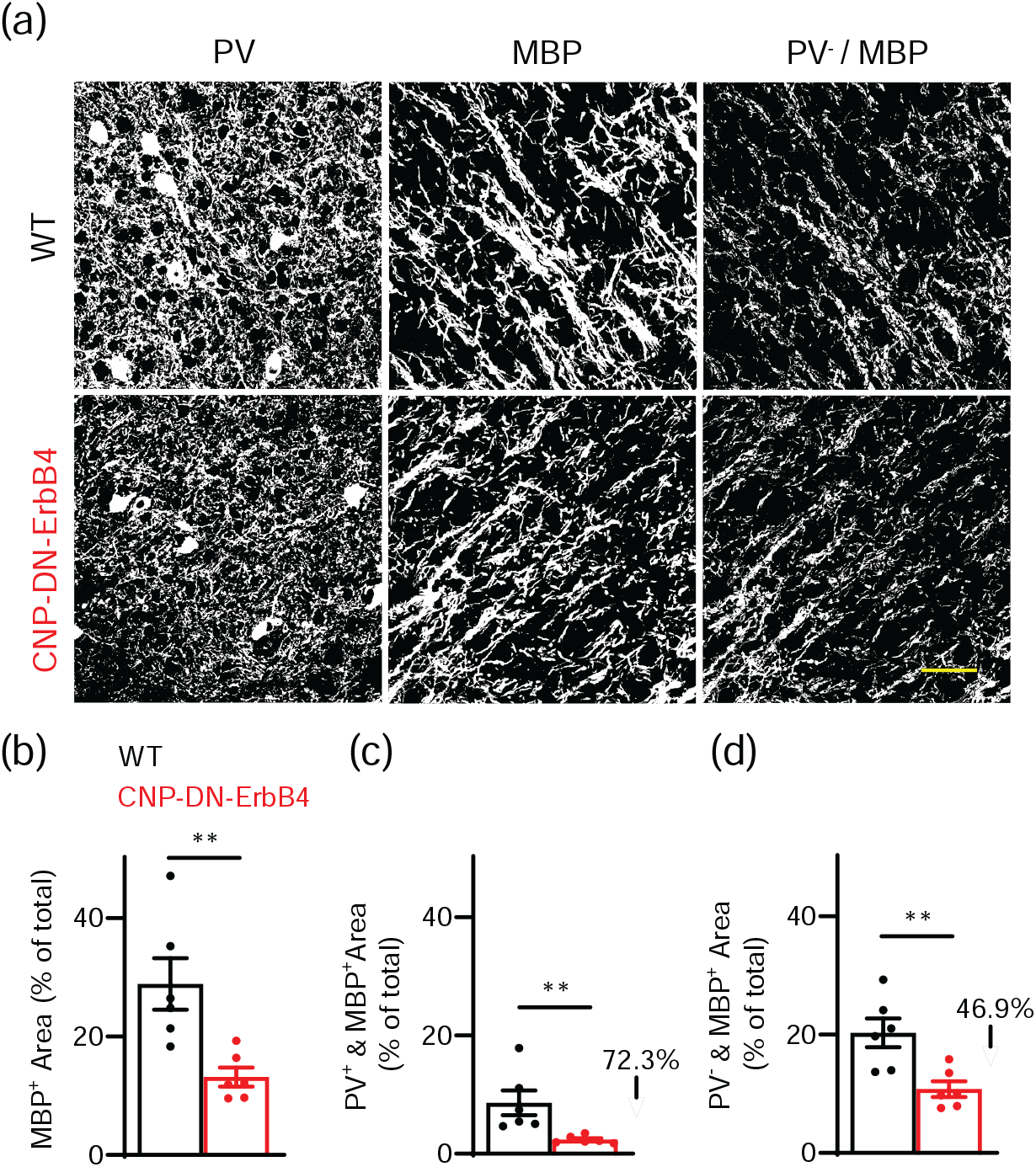
Loss of oligodendrocyte ErbB signaling leads to reduction in A1 MBP immunostaining both associated and not associated with PV immunostaining. **a**, Representative photomicrographs of A1 sections from WT and CNP-DN-ErbB4 mice showing the area with PV immunostaining (left), the area with MBP immunostaining (middle) and the MBP^+^ area not associated with PV staining (right). Scale bar: 20μm. **b**, Percent area covered by total MBP signal in A1 of WT (black) and CNP-DN-ErbB4 (red) mice (n = 6; p = 0.0043). **c**, Percent area covered by MBP signal associated with PV signal (PV^+^ & MBP^+^) in A1 of WT (black) and CNP-DN-ErbB4 (red) mice (n = 6; p = 0.0022). **d**, Percent area covered by MBP signal not associated with PV signal (PV^-^ & MBP^+^) in A1 of WT (black) and CNP-DN-ErbB4 (red) mice (n = 6; p = 0.0087). Mann-Whitney test was performed. Data are expressed as mean ± SEM.

### Oligodendrocyte-specific ErbB3 KO has the same effects on A1 myelination and PV^+^ neuron number as oligodendrocyte DN-ErbB4 expression

To test if the effects of hypomyelination on A1 interneurons occur in a different mouse model, we used a line in which we induce loss of oligodendrocyte ErbB signaling by a distinct approach i.e., inducible oligodendrocyte specific knockout of ErbB3, i.e. mice homozygous for an ErbB3 floxed allele (ErbB3^fl/fl^ (Qu et al., 2006)) and expressing the tamoxifen-inducible Cre recombinase (CreERT) under the control of the PLP1 promoter (PLP/creERT (Doerflinger, Macklin, & Popko, 2003)). We previously showed that these mice have CNS hypomyelination comparable to that seen in the CNP-DN-ErbB4 mice in other CNS regions (Makinodan et al., 2012), consistent with the overlapping expression of CNPase and Plp1 in mature oligodendrocytes (Marques et al., 2016).

Tamoxifen-induced oligodendrocyte ErbB3 KO led to a 28% reduction in A1 ErbB3 mRNA levels in the adult (Fig. 11a), consistent with loss of ErbB3 in oligodendrocytes, but not in the other cells expressing this receptor, including microglia, astrocytes, tanycytes, and some neurons (Gerecke et al., 2001). Since PV^+^ neurons do not ErbB3 receptors (Mayer et al., 2018; Que, Lukacsovich, Luo, & Foldy, 2021; Yau, Wang, Lai, & Liu, 2003) and Plp1 is expressed in myelinating cells (Marques et al., 2016), it is extremely unlikely that PV ErbB receptor signaling was altered in these mice. Oligodendrocyte ErbB3 KO led to a reduction in A1 MBP mRNA levels, indicative of hypomyelination (Fig. 11a). Like in CNP-DN-ErbB4 mice, oligodendrocyte ErbB3 KO also led to a reduction in A1 PV mRNA levels without affecting VIP and SST expression (Fig. 11b). In this model we again found a strong correlation between PV and MBP mRNA levels in A1 (r = 0.7787, p = 0.01; Fig. 11c). The number of PV^+^ cells in the A1 of PLP-ErbB3 KO mice was also significantly reduced respectively to controls (Fig. 11d,e).

**Figure 11:**
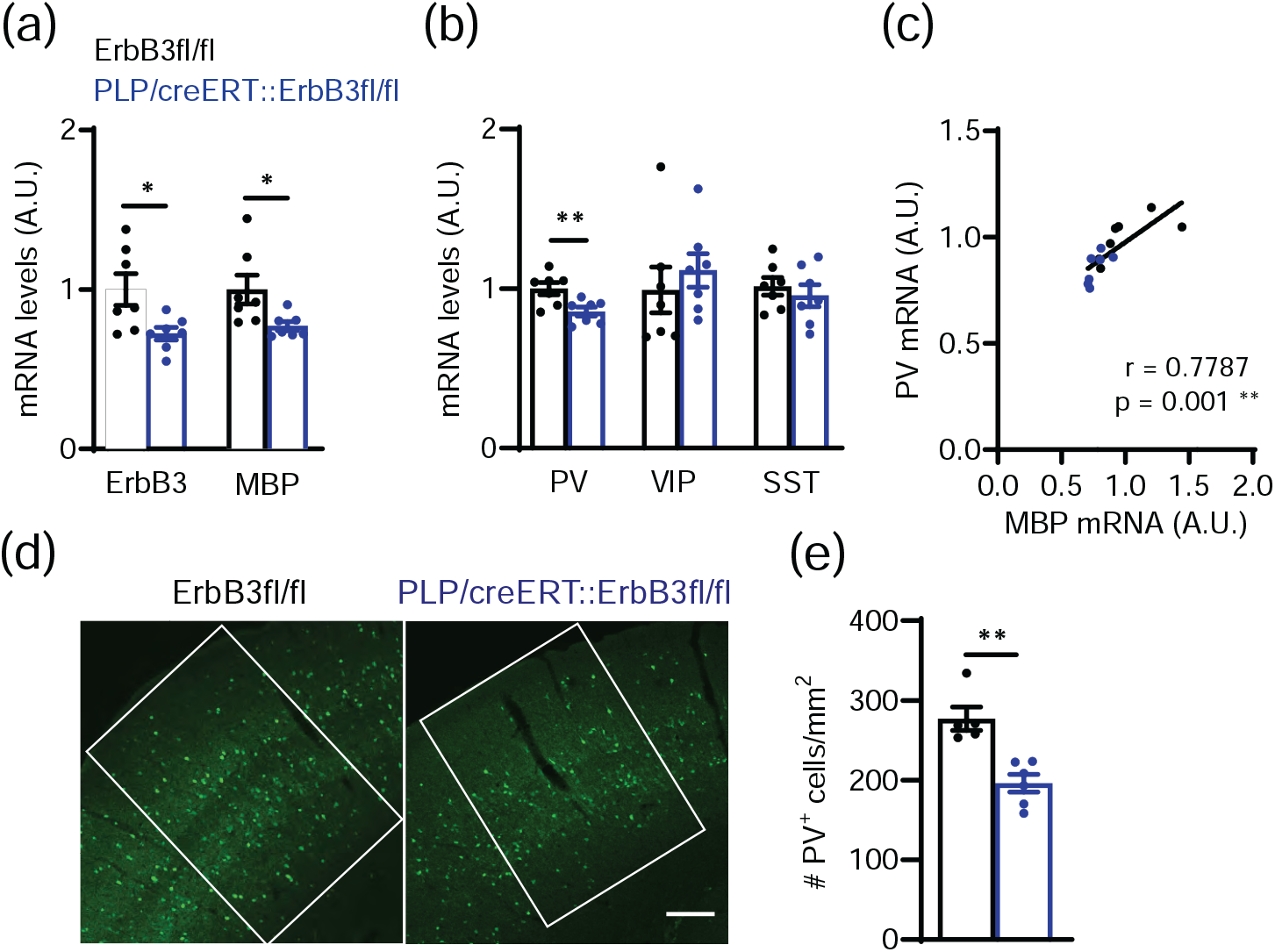
Loss of oligodendrocyte ErbB signaling by ErbB3 KO leads to reduced A1 MBP and PV mRNA levels and PV^+^ cell density. **a**, Reduced ErbB3 (n = 7; p = 0.0230) and MBP (n = 7; p = 0.0175) mRNA levels in A1 after oligodendrocyte-specific ErbB3 KO (PLP/creERT::ErbB3fl/fl). **b**, Relative mRNA expression for parvalbumin (PV), vasoactive intestinal peptide (VIP) and somatostatin (SST) in the A1 of WT (black) and PLP/creERT::ErbB3fl/fl (blue) mice. PV mRNA levels are reduced in A1 of PLP/creERT::ErbB3fl/fl mice (n = 6 – 7; p = 0.0093). No changes are observed in VIP (p = 0.3176) and STT (p = 0.62) mRNA expression. **c**, A1 PV mRNA levels correlate with MBP mRNA levels (WT: black dots, PLP/creERT::ErbB3fl/fl: blue dots; Pearson’s correlation r = 0.7787; p = 0.001). **d**, Representative photomicrographs of A1 sections from WT and PLP/creERT::ErbB3fl/fl mice showing PV^+^ cells (green). Scale bar: 100μm. **e**, A1 PV^+^ cell density is reduced in PLP/creERT::ErbB3fl/fl mice (n = 5 – 6; p = 0.0043). Mann-Whitney test was performed. Data are expressed as mean ± SEM.

Finally, to test if the impact of loss of oligodendrocyte ErbB receptor signaling on PV^+^ neurons is specific to A1, we quantified the number of PV^+^ cells in two other brain areas, the medial prefrontal cortex (PFC) and the primary motor cortex (M1), for both mutant lines and their respective wild type littermates. Consistent with the findings from A1, both CNP-DN-ErbB4 (Fig. 12a-d) and PLP-ErbB3 KO mice (Fig. 12e-h) showed reduced number of PV cells in the PFC and M1, compared to their littermate controls. These data indicate that hypomyelination leads to alterations in the number of PV-expressing neurons in the whole brain.

**Figure 12:**
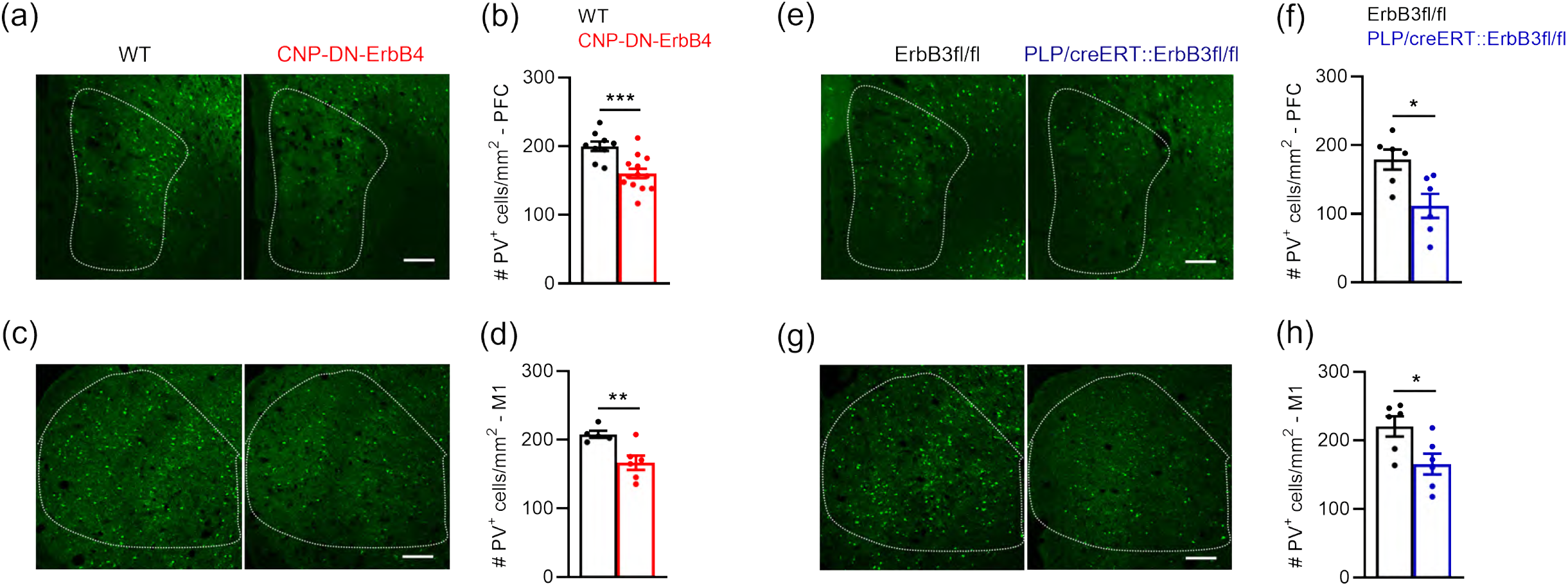
Loss of oligodendrocyte ErbB signaling by expression of DN-ErbB4 or ErbB3 KO leads to reduced PV cell density in other brain areas such as prefrontal cortex and primary motor cortex. **a**, Representative photomicrographs of prefrontal cortex (PFC) sections from WT and CNP-DN-ErbB4 mice showing PV^+^ cells (green). Scale bar: 100μm. **b**, PFC PV^+^ cell density is reduced in PFC of CNP-DN-ErbB4 mice (n = 9 – 13; p = 0.0009). **c**, Representative photomicrographs of primary motor cortex (M1) sections from WT and CNP-DN-ErbB4 mice showing PV^+^ cells (green). Scale bar: 100μm. **d**, PFC PV^+^ cell density is reduced in M1 of CNP-DN-ErbB4 mice (n = 5 – 6; p = 0.0083). **e**, Representative photomicrographs of prefrontal cortex (PFC) sections from WT and PLP/creERT::ErbB3fl/fl mice showing PV^+^ cells (green). Scale bar: 100μm. **f**, PFC PV^+^ cell density is reduced in PFC of PLP/creERT::ErbB3fl/fl mice (n = 6; p = 0.0411). **g**, Representative photomicrographs of primary motor cortex (M1) sections from WT and PLP/creERT::ErbB3fl/fl mice showing PV^+^ cells (green). Scale bar: 100μm. **h**, PFC PV^+^ cell density is reduced in M1 of PLP/creERT::ErbB3fl/fl (n = 6; p = 0.0260). Mann-Whitney test was performed. Data are expressed as mean ± SEM.

## DISCUSSION

Our results indicate that hypomyelination leads to altered excitatory/inhibitory balance in A1 cortical circuits due to diminished inhibitory connections to L2/3 neurons resulting in a shift towards excitation. These functional changes correlate with a reduced density of PV^+^ neurons, which, as illustrated by the preservation of VVA^+^ neuron density, reflects a loss of PV expression by some of the interneurons rather than the loss of the inhibitory neurons themselves. PV expression levels are regulated by neuronal activity (Dehorter et al., 2015; Favuzzi et al., 2017; Lagler et al., 2016; Stedehouder, Brizee, Shpak, & Kushner, 2018), and it has been proposed that myelin regulates PV^+^ neurons metabolism as well as improves the energy efficiency of signal propagation (Lee et al., 2012; Micheva et al., 2016; Rinholm et al., 2011). Thus, our results suggest that hypomyelination causes PV^+^ neuron hypoactivity, leading to the observed reduction in PV expression and inhibitory hypoconnectivity. These data, together with the tight correlation between MBP and PV expression, point to a powerful regulation of PV^+^ neuron function and gene expression by myelin. The observation that PV^+^ neuron density is also reduced in the prefrontal and motor cortices suggest that the impact of hypomyelination on PV^+^ neuron function is likely to occur in the whole brain.

The observation that hypomyelination alters excitatory/inhibitory balance in the cortex are similar to those made in mice with KO of the GABA_A_ receptor γ2 subunit in NG2+ cells, which include oligodendrocyte precursor cells (Benamer, Vidal, Balia, & Angulo, 2020). This study showed that barrel-cortex fast spiking interneurons in the GABA_A_R γ2 subunit KO mice present with severe myelin and axonal defects, as well as alterations in their function, leading to excitation-inhibition imbalance. Together, these observations suggest that communication between PV^+^ neurons and cells of the oligodendrocyte lineage are complex and could involve several signaling molecules and processes.

The finding that hypomyelination leads to reduced density of PV^+^ neurons but normal density of VVA^+^ cells is remarkably similar to what has been reported for several mutant mice that display autism spectrum disorder (ASD) endophenotypes, i.e., PV^+/-^, Shank1^-/-^ and Shank3B^-/-^ mice (Filice, Vorckel, Sungur, Wohr, & Schwaller, 2016). Interestingly, myelin defects have been found in ASD patients (Graciarena, Seiffe, Nait-Oumesmar, & Depino, 2018; Steinman & Mankuta, 2019), which also display auditory processing deficits (Jones et al., 2020; Rotschafer, 2021; Srinivasan, Udayakumar, & Anandan, 2020). Together, these observations raise the possibility that defects in myelination might contribute to some aspects of ASD by their negative impact on inhibitory networks.

The LSPS experiments show a significant spatial hypoconnectivity of inhibitory connections to L2/3 neurons in mice which hypomyelination, i.e., individual L2/3 neurons in the mutant mice receive inhibitory inputs from fewer locations, especially from L2/3 and L4. However, individual inhibitory connections are of comparable strength, indicating that hypomyelination causes a loss but not a weakening of connections. In contrast, excitatory connectivity in A1 appears to be normal. However, we cannot exclude the possibility that the effects of hypomyelination on excitatory axons are subtle and not resolved by our LSPS method. For example, hypomyelination could lead to increased conduction times in excitatory axons without loss of connections and cells, but since LSPS probes relatively short-range connections, such changes might be too small to be detected.

The specific effects of myelin on inhibitory circuits suggest that myelin might play multiple roles beyond controlling timing of axonal conduction. Our results suggest that hypomyelination affects axons of PV^+^ cells, which are fast-spiking neurons with metabolic demands higher than those of excitatory axons (Hu, Gan, & Jonas, 2014). In fact, the number of myelinated PV axons in A1 is reduced in the mutant mice, and the axonal PV fluorescence intensity and area strongly correlate with MBP fluorescence intensity and area. Of note, no alteration of expression of markers for two other important populations of inhibitory interneurons, the VIP and somatostatin, was found in A1 of mice with hypomyelination. This agrees with previous reports showing no myelin in VIP^+^ neurons and a rare myelination of somatostatin+ neurons in the somatosensory cortex (Micheva et al., 2016). Hypomyelination in both our mouse models affects the density of PV^+^ neurons in other brain regions, i.e., the prefrontal cortex and the primary motor cortex, supporting the notion that oligodendrocyte ErbB signaling is crucial for myelination of PV axons in the whole brain.

Our findings of altered circuits suggests that hypomyelination might lead to degraded fidelity of sound representation. This is consistent with the observation that focal demyelination of A1 L4 alters auditory frequency-specific responses (Narayanan et al., 2018) and abolishes tonotopic organization (Cerina et al., 2017). Since A1 neurons that encode the fast fluctuation of acoustic stimuli tend to receive balanced and concurrent excitation and inhibition (Bendor, 2015; Wang, Lu, Bendor, & Bartlett, 2008), our observation of altered inhibitory circuits in CNP-DN-ErbB4 mice are consistent with the functional deficits after hypomyelination. Thus, hypomyelination could play a role in the development of cognitive deficits. Indeed, we previously showed that CNP-DN-ErbB4 mice have behavioral phenotypes consistent with psychiatric disorders (Roy et al., 2007) and interneuron hypomyelination was reported in rat models of schizophrenia (Maas et al., 2020; Stedehouder & Kushner, 2017). Hence, our results showing reduced density of PV^+^ cells and altered network excitatory/inhibitory balance suggest that the absence of myelin might lead to circuit changes that manifest as impaired cognitive functions.

## Acknowledgements

This work was supported in part by NIH/NIDCD R01DC018500 (GC) and NIH/NIDCD R01DC009607 (POK).

## Data Availability

All data generated or analyzed during this study are included in the manuscript. Original blots and images used for calculations, Matlab codes, and maps into 3D matrices used in electrophysiology experiments will be uploaded to Dryad.

